# Who reviews for predatory journals? A study on reviewer characteristics

**DOI:** 10.1101/2020.03.09.983155

**Authors:** Anna Severin, Michaela Strinzel, Matthias Egger, Marc Domingo, Tiago Barros

**Affiliations:** Doctoral researcher at the Swiss National Science Foundation (SNSF) and the University of Bern, Switzerland; Scientific collaborator at the SNSF, Bern, Switzerland; President of the National Research Council of the SNSF and professor at the University of Bern, Switzerland; Product development intern at Publons, London, UK; Product lead at Publons, London, UK

## Abstract

**Background:** While the characteristics of scholars who publish in predatory journals are relatively well-understood, nothing is known about the scholars who review for these journals. We aimed to answer the following questions: Can we observe patterns of reviewer characteristics for scholars who review for predatory journals and for legitimate journals? Second, how are reviews for potentially predatory journals distributed globally?

**Methods:** We matched random samples of 1,000 predatory journals and 1,000 legitimate journals of the Cabells Scholarly Analytics’ journal lists with the Publons database of review reports, using the Jaro-Winkler string metric. For reviewers of matched reviews, we descriptively analysed meta-data on reviewing and publishing behaviour.

**Results:** We matched 183,743 unique Publons reviews that were claimed by 19,598 reviewers. 6,077 reviews were conducted for 1160 unique predatory journals (3.31% of all reviews). 177,666 were claimed for 6,403 legitimate journals (96.69% of all reviews). The vast majority of scholars either never or only occasionally submitted reviews for predatory journals to Publons (89.96% and 7.55% of all reviewers, respectively). Smaller numbers of scholars claimed reviews predominantly or exclusively for predatory journals (0.26% and 0.35% of all reviewers, respectively). The two latter groups of scholars are of younger academic age and have fewer publications and fewer reviews than the first two groups of scholars.Developing regions feature larger shares of reviews for predatory reviews than developed regions.

**Conclusion:** The characteristics of scholars who review for potentially predatory journals resemble those of authors who publish their work in these outlets. In order to combat potentially predatory journals, stakeholders will need to adopt a holistic approach that takes into account the entire research workflow.

## Introduction

Scholars spend a considerable amount of their time reviewing the work of their peers. A recent study suggests that 13.7 million reviews are done every year and that writing a review takes a median five hours [1]. When assessing a manuscript for its soundness, originality, validity, and possible impact, reviewers serve a critical gatekeeping function: they help ensure that only papers, which pass a certain quality threshold, enter the scholarly record [2,3]. Such gatekeeping is particularly important in fields where practitioners and policymakers rely on evidence in the form of published journal articles. Without rigorous peer review, invalid studies could influence policies and practices and potentially cause harm to the population [4].

Reviewing for so-called predatory journals might, however, be a waste of valuable time and effort. Prioritizing self-interests at the expense of scholarship, these outlets exploit the open access (OA) model of publishing. They offer to publish articles but do not provide the services one would expect to receive from a legitimate journal [5–7]. Amongst other deficiencies, predatory journals do not guarantee archiving and long-term access to their contents [6,8]. For this reason, their publications generate limited readership, are rarely cited, and might eventually be lost [9]. Reviewing a manuscript that cannot be found, and will hardly produce any scientific impact, is unlikely to advance science. Also, a researcher who submitted purposively flawed manuscripts to predatory journals, showed that where reviewers spotted severe scientific problems, editors accepted some papers nevertheless [10]. Just as publishing research in predatory journals, reviewing for these outlets can be described as a waste of vital resources and should, therefore, be stopped [11].

While the profiles of scholars who publish in predatory journals and who cite these articles are well-studied [11–16], little is known about the scholars who contribute to the peer review of these outlets. Amongst other factors, this is because peer review, like most other forms of academic gate-keeping, often is not publicly observable [17]. This is particularly true for predatory journals, which lack transparency of publishing processes and policies [18]. Understanding who reviews for predatory journals might help to gain insights into the peer review processes of these outlets and thus assist in combating them. We aimed to analyze the characteristics of scholars who have claimed reviews on Publons reviews for predatory journals and to legitimate journals, as defined by Cabell’s Scholarly Analytics’ journal lists. We analyzed their sociodemographic characteristics as well as their reviewing and publishing behavior.

## Material and Methods

To study characteristics of scholars who reviewed for predatory journals and legitimate journals, we matched random samples of journals indexed in Cabell’s Scholarly Analytics’ (hereafter Cabell’s) journal lists with the Publons database of peer review reports. For the scholars who provided matched reviews, we analyzed publishing and reviewing behavior and sociodemographic characteristics through descriptive statistics.

### Data sources

As the basis for identifying reviews for predatory journals and likely legitimate journals, we used Cabell’s Scholarly Analytics’ (hereafter Cabell’s) journal lists. Cabell’s is a scholarly services company that maintains a list of likely legitimate journals and a list of journals “as potentially not following scientific publication standards or expectations on quality, peer reviewing, or proper metrics” [19]. To be indexed in either list, a journal must fulfill several requirements that cover different aspects, including business practices, editorial services, publishing policies, and archiving and access [7]. Both Cabell’s lists are subscription-based. We purchased access and downloaded the list of predatory journals and the list of legitimate journals in December 2018. The lists contained 10,671 and 11,057 unique journal titles, respectively. Of note, lists of potentially predatory and legitimate journals are inconsistent and sometimes out of reach [5]. In addition, predation in academic publishing is not a simple binary phenomenon but should be understood as a spectrum with varying degrees of illegitimacy [17]. Some journals that have been classified as either predatory or legitimate might actually operate in a grey zone.

Publons is a platform for researchers to track their scholarly contributions and get recognition for their peer reviews. Researchers sometimes provide Publons evidence of reviews that they performed for predatory journals. Publons does not endorse these journals but chooses to display the reviews to provide greater transparency into the peer review practices and communities surrounding these journals. Further information about Publons’ approach displaying review and journal information is available from Publons’ guidelines to evaluating publishers [20]. As of October 2019, Publons contained data on more than 5 million reviews, spanning more than 500,000 reviewers and approximately 40,000 journals [21].

### Study sample

In a first step, we drew random samples of 1,000 predatory journal titles and 1,000 legitimate journal titles of the Cabell’s journal lists. In a second step, we matched both samples with the Publons entire database of review reports, to identify reviews for predatory journals and reviews for legitimate journals. Following an approach used by Strinzel et al. (2019), this matching was based on journal names [7]. Because of potential typos and orthographical differences between the Cabell’s lists and the Publons database, we matched journal names based on their relative similarity, using the Jaro-Winkler algorithm in R package RecordLinkage [7,22]. The Jaro-Winkler metric, which ranges from 0 (no similarity) to 1 (exact match), was calculated for all possible pairs of journals [23]. Having calculated the Jaro-Winkler metric, we manually inspected all pairs with a Jaro-Winkler metric smaller than 1 to identify pairs where due to orthographical differences between the lists, no exact match was found. For each matched pair, we further compared the journals’ publishers to exclude cases where two journals had the same or a very similar name but were, in fact, two different outlets.

Only journals for which we found at least one review in the Publons database were analyzed further. In the third step, we created an extended network of journals. For scholars who we found to have reviewed for matched journals, we selected all other journals for which they also claimed one or more reviews on Publons. In a fourth step, we matched all journals of the extended network against the Cabell’s lists, to classify their reviews as for either predatory or legitimate journals. Doing so, we again calculated the Jaro-Winkler metric and did a manual assessment of all journal pairs, as done in the second step. All steps taken in identifying scholars who reviewed for predatory journals and legitimate journals are shown in Figure 1. For reviewers of matched reviews, we retrieved data on their sociodemographic characteristics and publishing and reviewing behavior from the Publons database. The data covered their primary institutional affiliation, dates of first publication and last publication, number of publications, and number of reviews in Publons database. Based on these variables, we calculated proxies for academic age (based on the date of the first and last publication) as well as for publishing and reviewing productivity (as the average number of articles published and reviews done per year, respectively). We further matched the country of the primary institutional affiliation with World Bank data [24], to classify countries according to their region and income. We analyzed reviewer characteristics through descriptive statistics.

**Figure 1:**
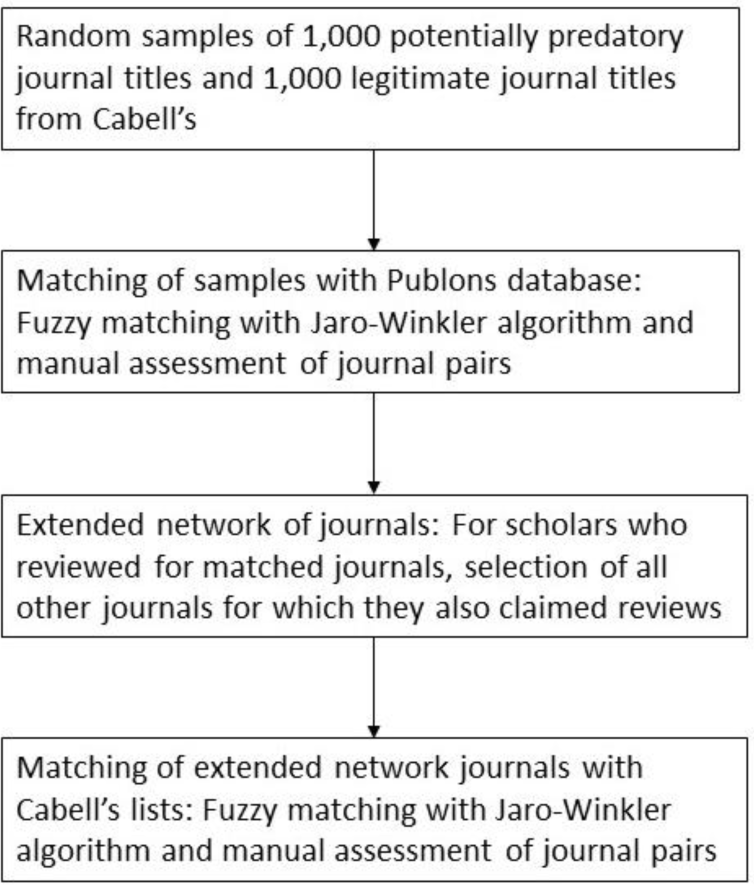
Procedure of identifying reviewers for predatory journals and legitimate journals.

## Results

The sampling and matching procedure resulted in 183,743 unique Publons reviews that were submitted by 19,598 reviewers. A total of 6,077 reviews were done for 1160 unique predatory journals (3.3% of all reviews of the matched journals), and 177,666 for 6,403 legitimate journals (96.7% of all reviews of the matched journals).

### Sociodemographic characteristics and publishing and reviewing behavior

We analyzed peer-reviewing activities and reviewer characteristics based on the retrieved meta-data on publishing and reviewing behavior (i.e., number of publications and reviews, dates of first and last publication). This information was available for 86,298 unique reviews submitted to Publons by 7349 individual reviewers. Based on each reviewer’s share of reviews for predatory journals, we created five subgroups of reviewers (Table 1). Most reviewers had never submitted a review for a predatory journal to Publons (90% of all reviewers). This group had a relatively old academic age, was well published but had submitted relatively few reviews to Publons. A small group of reviewers (8%) had occasionally reviewed for predatory journals, with shares of predatory reviews ranging from one to 25% of all review. Reviewers in this group were experienced and productive: they had the oldest academic age, published the largest number of articles and submitted the second-highest number of reviews on Publons (Table 1). Only about two percent reviewed regularly for predatory journals (share 26% to 75%) and they occupied a intermediate position in terms of academic age and publication and review activity. Very few scholars had shares of predatory reviews between 76 and 99% (0.26% of all reviewers). Interestingly, they had the youngest academic age and lower number of publications than the first three groups, but submitted a high number of reviews to Publons. A similarly small number (0.35% of all reviewers) exclusively reviewed for predatory journals. They had a young academic age, published few articles, and submitted few reviews to Publons.

**Table 1:** Reviewer subgroups and their characteristics.

| Subgroup, based on shares of predatory reviews per reviewer | No. of reviewers (share of all 7349 reviewers) | Mean academic age (SD) | Mean no. of reviews per reviewer (SD) | Mean no. of publications per reviewer (SD) | Reviews written per year per reviewer (SD) | Articles published per year per reviewer (SD) |
| --- | --- | --- | --- | --- | --- | --- |
| **No predatory reviews (0%)** | 6611 (90.0%) | 15.27 (10.28) | 46 (77.85) | 57.28 (73.71) | 3.65 (6.81) | 3.25 (3.68) |
| **Few predatory reviews (1-25%)** | 555 (7.6%) | 17.37 (11.39) | 132.3 (153.63) | 85.39 (119.08) | 9.9 (19.27) | 4.43 (3.87) |
| **Some predatory reviews (26-75%)** | 138 (1.9%) | 12.47 (10.42) | 107.8 (173.96) | 49.58 (75.26) | 10.60 (19.64) | 3.42 (3.33) |
| **Many predatory reviews (76-99%)** | 19 (0.26%) | 8.16 (6.53) | 236.6 (401.20) | 43.53 (75.74) | 36.71 (60.29) | 5.29 (12.9) |
| **Predatory reviews only (100%)** | 26 (0.35%) | 9.38 (6.27) | 26.35 (40.82) | 18 (14.49) | 3.34 (5.64) | 1.91 (1.21) |

### Geographic distribution of reviews

; Based on the reviewer’s primary institutional affiliation, we analyzed the geographic distribution of reviews. This information was available for 10,950 reviewers affiliated with institutions in 117 different countries, who submitted 107,722 reviews. To interpret reviewing behavior in the context of reviewing productivity, we calculated shares of reviews for predatory journals and shares of reviews for legitimate journals for each region and World Bank country income group. Regions with the highest shares of predatory reviews were Sub-Saharan Africa (21.8%), Middle East and North Africa (13.9%) and South Asia (7.0%), followed by North America (2.1%), Latin America and the Caribbean (2.1%), Europe and Central Asia (1.9%) and East Asia and the Pacific (1.5%) (Figure 2). Low-income countries had the highest share of predatory reviews (27.0%), followed by lower-middle-income countries (17.5%), upper-middle-income countries (4.9%) and high-income countries (1.5%) (Figure 3).

**Figure 2:**
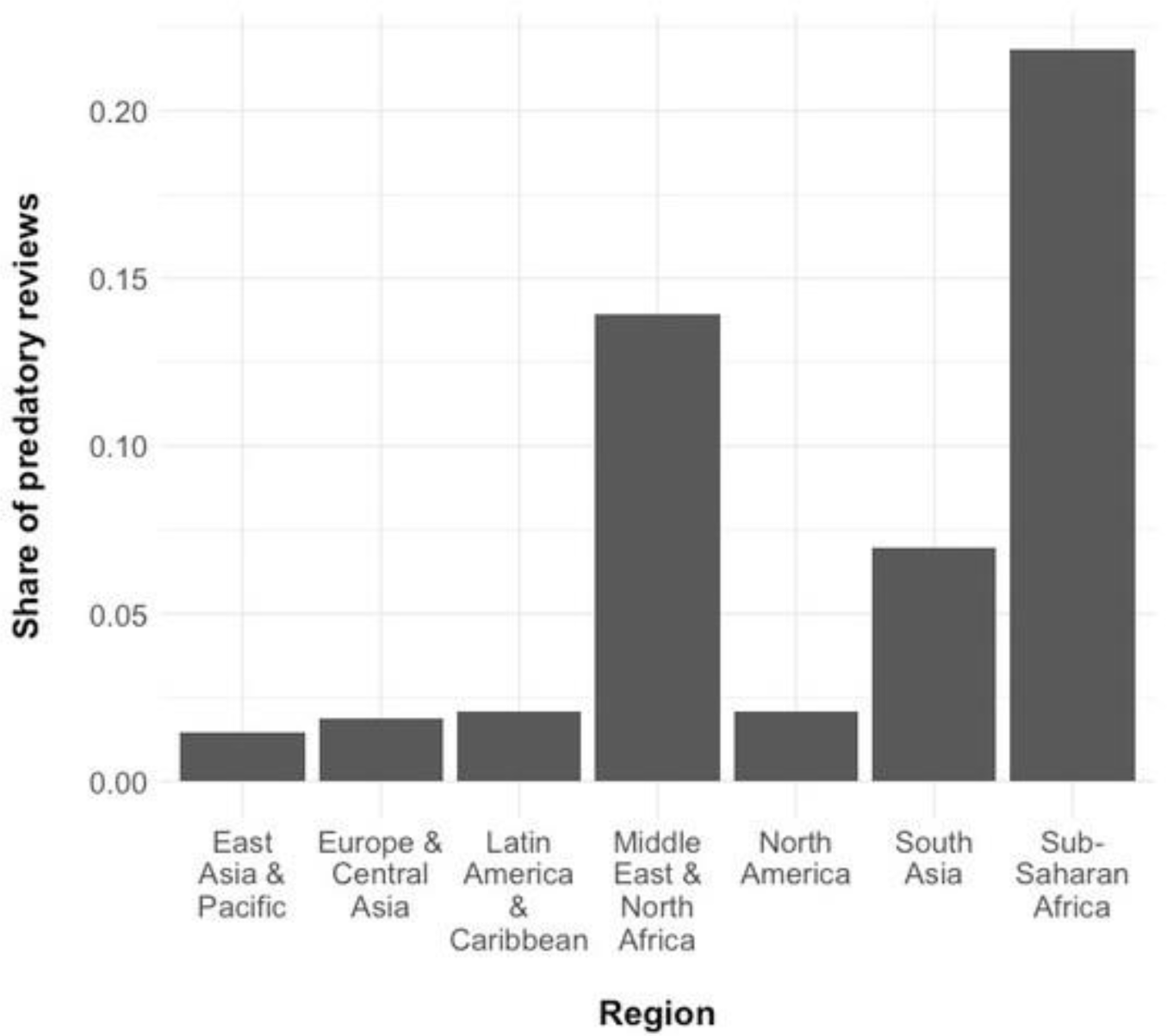
Shares of predatory reviews by region.

**Figure 3:**
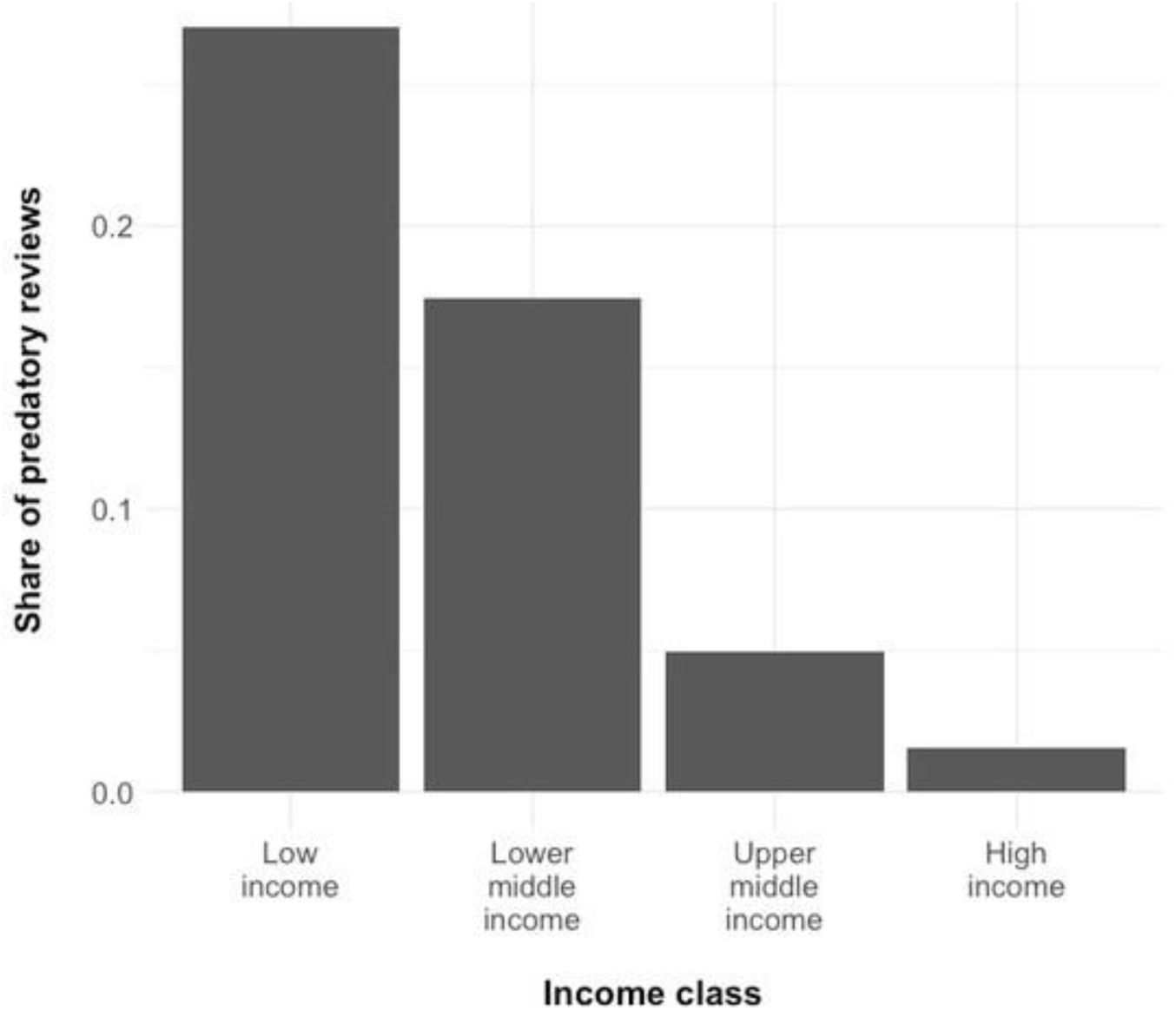
Shares of predatory reviews by World Bank income class.

## Discussion

This study investigated the profiles of scholars who review for predatory journals and legitimate journals. It examined the reviewing patterns for different subgroups of scholars as well as their publishing behavior, scientific productivity, and academic age. This study further investigated the geographic distribution of reviews for predatory journals and legitimate journals.

Matching journal titles from the Cabell’s list of predatory journals and the Cabell’s list of legitimate journals with the Publons database of peer review reports resulted in almost 200,000 unique reviews that were submitted by nearly 20,000 reviewers. Out of these, approximately 6,000 reviews were written for predatory journals, indicating that some predatory journals do conduct some form of peer review, or at least solicit it. Even though the number of predatory reviews was relatively small, it still corresponds to a significant expenditure of time. According to recent estimates [1], it will have taken approximately 30,000 hours to write these reviews – time that scholars wasted on manuscripts that will generate limited scientific impact.

The analysis of reviewing patterns showed that scholars who reviewed predominantly or exclusively for predatory journals appear to be less experienced than reviewers with no or only a few predatory reviews: they had a younger academic age, fewer publications, and wrote fewer reviews. Their characteristics resemble those of authors publishing in predatory journals, who are inexperienced and young scholars [12]. The geographical distribution of reviews submitted for predatory journals showed that low-income and lower-middle-income countries have particularly high shares of predatory reviews submitted to Publons. In contrast, upper-middle-income and high-income countries show very low shares of predatory reviews. A possible explanation for this observation is that predatory journals have become an integral part of the workflow for many scholars in low- and lower-middle income countries. Putting our results in context with the literature [10,12,15,25], we believe that predatory journals are relevant for scholars in these regions not only for publishing their work but also for reviewing the work of others. When tracing the location of journal editors, Bohannon found that many of the journals accepting his intentionally flawed manuscript were based in developing countries (particularly India and Nigeria) with branches in the USA and the UK [10]. Also, the authors of publications in predatory journals predominantly are located in developing countries. Researchers who published in predatory journals were mainly from developing regions, particularly South Asia (primarily India and Pakistan) and Africa (mainly Nigeria). For researchers in North America, Australia, and Europe, predatory journals were less relevant [12].

Inexperienced scholars and scholars in developing countries might be more likely to be tricked into believing that they review for a legitimate journal. It is also possible that predatory journals provide an opportunity for marginalized members of the global academic community to survive in the “publish or perish” culture of today’s academia. Traditionally, a scholar’s track record or the ability to secure funding outweigh peer review contributions when it comes to academic promotion or funding decisions [26]. Nevertheless, scholars now increasingly make use of platforms that help them record, verify, and showcase their reviewing activities on their applications or their CVs [27,28]. In an academic system that is characterized by a pressure to be productive and distinguish oneself [29], particularly early-career researchers and researchers from developing countries might be eager to expand the list of journals for which they reviewed.

This study suffers from a few limitations. First, whether a review was written for a predatory journal or a legitimate journal does not allow us to draw any conclusions on its quality or integrity. Our study was not designed to evaluate the quality of individual reviews or whether editors followed the recommendation of reviewers. Some experiments, however, showed that peer review in predatory journals was superficial and focused on a manuscript’s layout rather than on its scientific problems [10,17]. Also, where reviewers spotted severe scientific problems, editors accepted some papers nevertheless [10]. In a separate study, we, therefore, are investigating whether the quality of peer review differs across different types of journals, including predatory journals and legitimate journals. Second, lists of journals are difficult to keep up do date and might not reflect recent changes in publishing practices of journals. Cabell’s lists have further been shown to neglect important but challenging to verify criteria, including peer review and editorial services [7]. We hence might have included false positives and excluded false negatives from our analysis. Some journals might also operate in a grey zone and cannot be easily classified as either predatory or legitimate [4,7]. Third, we did not analyze data on when a review was submitted to Publons, which limited our ability to draw conclusions on the correlation between academic age and reviewing behavior. Finally, there might be self-selection biases present in submitting reviews to Publons. For example, reviewers might be more likely to provide reviews for legitimate or high-quality journals to Publons than for predatory or low-quality journals. Such bias will probably have led us to underestimate shares of predatory reviews.

Most initiatives dealing with the problem of predatory journals focus on reducing the submissions of manuscripts by warning authors not to publish their work in these outlets [4]. In order to prevent scholars from reviewing for predatory journals, research institutions, funders, and publishers ought to increase the training of reviewers. Besides teaching scholars how to conduct a review, this should involve enabling them to make an informed decision for which journals they review. A useful resource for this could be an adapted version of the initiative *<u>Think, Check, Submit</u>*. Funders and research institutions might further monitor for which journals their grantees and faculty members review and warn against using paid research time to review for predatory journals. When evaluating applications for funding or promotion, they might check peer review records for predatory titles. Services that help researchers get credit for their reviewing activities should have a clear policy for predatory journals. Because lists of predatory journals and legitimate journals tend to lack transparency, services that check peer-reviewing activities might, therefore, assess the contents of submitted reviews for quality and rigor. Research institutions, funders, and publishers should boost their efforts in discouraging scholars from reviewing for these outlets.

To our knowledge, this is the first study that investigates the profiles of scholars who review for predatory journals. Our study suggests that there are fw reviews submitted to Publons for predatory journals but that writing these reviews still meant a considerable expenditure of time and resources. Our study further finds that the profiles of scholars who review for predatory journals tend to resemble those scholars who publish their research in these outlets: they tend to be young and inexperienced researchers who are affiliated with institutions in developing regions. Because predatory journals have gained relevance for scholars not only for publishing their own work but also for reviewing the work of other researchers, a holistic approach to combating these journals is needed that takes into account the entire research workflow. More research is necessary that investigates the quality of peer review across different types of journals, including potentially predatory journals and legitimate journals.

